# SMITH: Spatially Constrained Stochastic Model for Simulation of Intra-Tumour Heterogeneity

**DOI:** 10.1101/2022.07.22.501136

**Authors:** Adam Streck, Tom Kaufmann, Roland F. Schwarz

**Author notes:** These authors contributed equally (Adam Streck, Tom Kaufmann).

## Abstract

**Motivation:** Simulations of cancer evolution and cellular growth have proven highly useful to study, in detail, the various aspects of intra-tumour heterogeneity, including the effect of selection, mutation rates, and spatial constraints. However, most methods are computationally expensive lattice-embedded models which cannot simulate tumours with a realistic number of cells and rely on various simplifications. Alternatively, well-mixed stochastic models, while efficient and scalable, do not typically include spatial constraints and cannot reproduce the rich clonal dynamics observed in real-world tumours.

**Results:** We present SMITH, a simple, efficient, and explainable model of cancer evolution that combines the advantages of well-mixed stochastic models with a new confinement mechanism which limits the growth of clones based on the overall tumour size. We demonstrate that this confinement mechanism is sufficient to induce the rich clonal dynamics observed in spatial models, while allowing for a clear geometric interpretation and efficient simulation of one billion cells within a few minutes on a desktop PC. We explore the extent of stochasticity and rigorously assess the effects of cell turnover, mutation rate, fitness effects and confinement on the resulting clonal structures.

**Availability and Implementation:** SMITH is implemented in C# and freely available at bitbucket.org/schwarzlab/smith together with binaries for all major platforms. For rich visualisations of the simulated clonal dynamics we provide an accompanying Python package PyFish at bitbucket.org/schwarzlab/pyfish.

**Supplementary information:** All supplementary figures are in the supplementary document.

## 1 Introduction

Carcinogenesis is governed by random mutational processes influenced by cell-intrinsic and environmental factors and shaped by selection, which result in the successive accumulation of genetic aberrations. Despite the randomness of the underlying processes, cancers ultimately converge towards a common set of phenotypic traits, such as replicative immortality, immune evasion, and invasiveness, commonly known as the Hallmarks of Cancer (Hanahan, 2022). The progenies of the tumour-originating cell continue to accumulate mutations past malignant transformation, and this continuous evolution and selection (Watkins *et al*., 2020) gives rise to significant genomic differences between clones inside a tumour, known as *intra-tumour heterogeneity* (ITH). ITH has been linked to progression, metastasis, treatment resistance and overall poor patient outcomes (Marusyk *et al*., 2012; Beerenwinkel *et al*., 2015; Jamal-Hanjani *et al*., 2017; Turajlic *et al*., 2018; Watkins *et al*., 2020). Understanding the aetiology of ITH is thus key to successful treatment of cancer and to prevent resistance development (Marusyk *et al*., 2020).

To study ITH and cancer evolution, two orthogonal approaches are common. Most studies focus on retrospective inference of the evolutionary history of a tumour from e.g. sequencing data of clinical tumour specimens. Such retrospective studies have accurately described the mutational processes shaping cancer genomes and greatly improved our understanding of cancer evolution (Steele *et al*., 2022; Drews *et al*., 2022; Watkins *et al*., 2020; Turajlic *et al*., 2018; Jamal-Hanjani *et al*., 2017). Alternatively, forward simulations of cancer evolution allow direct testing of different biological hypotheses and modelling assumptions, which can then be contrasted with empirically observed patterns of ITH (Beerenwinkel *et al*., 2015). Due to computational advances and the increasing availability of data, such forward simulations play an increasingly important role (Noble *et al*., 2022; West *et al*., 2021; Zhao *et al*., 2021; Watson *et al*., 2020; Chkhaidze *et al*., 2019).

Many computational models have been proposed for simulating cancer evolution and these models often employ variants of cellular automata, where cells or groups of cells are positioned on a 2D or a 3D lattice (Iwasaki and Innan, 2017). The lattice embedding has the advantage that it directly creates spatial constraints which enable the simulation of e.g. biopsy results (Chkhaidze *et al*., 2019) or the dispersal of cells in space (Waclaw *et al*., 2015), or between neighbouring tissues (Noble *et al*., 2022). However, the lattice structure also induces limitations, for example by fixing the number of neighbouring cells in 3D to either 6, 14, or 26 cells for the von Neumann, Hexagonal, or Moore neighbourhoods respectively (Iwasaki and Innan, 2017). In addition, competition for space is difficult to model and computationally complex, and simplifying rules are frequently employed, such that a whole row of cells needs to be moved at once (Chkhaidze *et al*., 2019), or that dead cells have to disappear from the lattice (West *et al*., 2021). These rules often do not reflect actual cell mechanisms and are mainly driven by complexity constraints. Despite these optimisations, simulating individual cells to the size of a real tumour of 1-2 cm in diameter (Erdi, 2012) comprising around 1 billion cells (Alberts *et al*., 2002; Del Monte, 2009) remains difficult even on supercomputer architecture (Rosenbauer *et al*., 2020). Thus, cells are usually grouped to uniform populations of glands (Sottoriva *et al*., 2015), demes (Noble *et al*., 2022), or severely limited in size (West *et al*., 2021).

Conversely, stochastic models of well-mixed populations, such as the commonly used branching process model of cancer (Haccou *et al*., 2005), are highly scalable, but assume an exponentially growing population without spatial constraints. These unconstrained models only exhibit a limited amount of clonal dynamics and are characterised by a low number of driver mutations and low to medium clonal diversity (Noble *et al*., 2022). They have been successfully applied to modelling clonal haematopoesis, where space is not a primary limiting factor (Watson *et al*., 2020), but their applicability to solid tumours remains limited. Therefore there is a general need for efficient and scalable models of tumour evolution that include spatial constraints.

To address the above listed issues, we introduce SMITH (Stochastic Model of Intra-Tumour Heterogeneity), a fast stochastic model of cancer evolution with spatial constraints that can simulate realistically sized tumours of up to one billion cells. At its base, SMITH employs a classical branching model of cancer, where cell birth and death are driven by random processes (Haccou *et al*., 2005), modulated by fitness-increasing mutations, and without an explicit representation of cell location. Additionally, SMITH introduces the concept of confinement, a simple mechanism that limits the growth and turnover of a population of cells based on the size of that population. We formulate confinement in terms of the 3D geometry of a tumour, where it serves to separate the tumour into a proliferating shell and a static core. The size of the shell can be adjusted, and confinement thus provides a natural way to embed both purely well-mixed and surface-growth-only tumour models into simulations of cancer evolution, without keeping track of the location of individual cells or clones. Using SMITH we demonstrate that confinement alone is sufficient to reproduce clonal dynamics typical for explicitly spatial simulations and spatially organised tumours. We explore the dynamics of fitness generating and accumulating functions and investigate their effect on cell birth and death and the clonal dynamics of the tumour. Using repeated simulations of tumour growth with one billion cells we investigate the effect of randomness on these clonal dynamics and show that even a simple model with limited stochasticity leads to a wide variety of possible clonal architectures and dynamics.

## 2 Methods

The SMITH model stochastically describes the size of tumour cell populations over time, thereby keeping track of the number of cells, any mutations they might have acquired and the evolutionary relationship between them. SMITH is based on a Galton-Watson branching process (Roch, 2015) and makes use of four key assumptions: (i) a low mutation probability with a non-negative fitness effect (driver mutations), (ii) a well-mixed population of cells, (iii) a spherical tumour shape, and (iv) confined growth. Assumptions (i) and (ii) are common in branching process modelling, and (iii) describes a well-studied group of tumours (Black and McGranahan, 2021). The assumption (iv) is specific to our model and directly changes the dynamics of the system. Under confinement, we split the tumour into two distinct regions: the core which contains “confined” cells that cannot undergo cell division and the outer layer, termed “shell”, in which the cell turnover takes place.

### 2.1 Model overview

A tumour as modelled by SMITH comprises three different types of cells: *alive* cells, *removed* cells, and *necrotic* cells. A newly created cell is always *alive*. Upon its death it can either be degraded (*removed*) or remains embedded in the tumour as a *necrotic* cell, where it continues to contribute to the overall tumour mass, but does not divide any longer.

Since SMITH does not model cellular position explicitly, it only keeps track of the number of cells in clones, where each clone is a collection of cells sharing the same set of mutations. Formally, a clone is a 4-tuple *c* = (*c*_*a*_, *c*_*r*_, *c*_*n*_, *c*_*M*_), where *c*_*a*_, *c*_*r*_, *c*_*n*_ ∈ ℕ_0_ describe the number of alive, removed, and necrotic cells in that clone respectively and *c*_*M*_ describes the set of mutations shared by all the cells in *c*. Each mutation from the set of all possible mutations (*m ∈ ℳ*) is defined by a unique identifier. The set of possible mutations *ℳ* is thereby considered infinite (infinite sites assumption (Kimura, 1969)), so that every mutation *m* can only occur once.

The model is parameterised by the parameter vector *θ* = (*θ*_turn_, *θ*_mut_, *θ*_fit_, *θ*_conf_) ∈ [0, 1]^4^, which describes the cell turnover probability, the mutation probability, the average fitness increase of a mutation, and the confinement value, and which are described in their corresponding sections below. The SMITH model is further configured by three different modelling options (Methods Sec. 2.4): the fitness distribution *o*_dis_ from which fitness values of new mutations are drawn, the mutation effect *o*_eff_ which determines if a mutation affects cell birth, death, or both, and the accumulation method *o*_acc_ which defines how the fitness values of individual mutations in a clone are aggregated to form the clone-level fitness value.

To describe the state of the system over time *t ∈* ℕ_0_, we denote as *C*^*t*^ the set of clones at time step *t*. All simulations start from a single clone with a single alive cell characterised by a single identifying mutation, i.e. *C*^0^ = {(1, 0, 0, {*m*})}. A simulation is then a transformation of *C*^*t*^ into *C*^*t*+1^ under the parameter set *θ* and with the model options *o*. For brevity we sometimes use counts for alive, removed and necrotic cells across the whole population *C*^*t*^ via 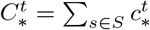 for the time *t*.

We use two stopping conditions for the simulation: the maximum number of time steps *max steps* and the maximum population size *max pop*. The simulation stops at a step *t* when *t* = *max steps* or 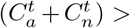 *max pop*. We also require the model to reach a minimum population *min pop*. If the simulation terminates while 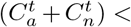 *min pop* and *t < max steps*, the result is discarded and the simulation restarts. All the parameters, options, and conditions can be found in Table 1.

**Table 1.** Overview of all relevant model parameters and variables. Note that only the parameters affect the behaviour of the model. The variables only control the length of the execution and the output.

| parameters |  | options |  | conditions |  |
| --- | --- | --- | --- | --- | --- |
| $\theta_{\text{turn}}$ | 0.01 | $o_{\text{dis}}$ | *const* | *min_pop* | 1000 |
| $\theta_{\text{conf}}$ | 0.1 | $o_{\text{acc}}$ | *mul* | *max_pop* | *min_pop* · 2<sup>20</sup> |
| $\theta_{\text{mut}}$ | 10<sup>-4</sup> | $o_{\text{eff}}$ | *birth* | *max_steps* | 10<sup>6</sup> |
| $\theta_{\text{fit}}$ | 0.2 | | | *cutoff* | 0.001 |

### 2.2 Cell turnover

We start with a basic system in homeostasis where the number of cells is kept constant on average and without novel mutations or necrosis. We thus define a cell birth and death process by sampling the number of cells that are born *B*_*b*_(*c*^*t*^) and that have died *B*_*d*_(*c*^*t*^) at time step *t* in clone *c* from a binomial distribution with the birth probability *p*_birth_ = *θ*_turn_ and the death probability *p*_death_ = *θ*_turn_. The new number of alive and removed cells at time step *t* + 1 is then defined as:

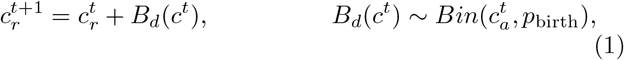

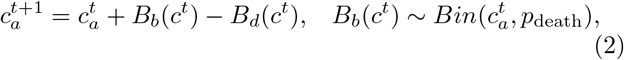

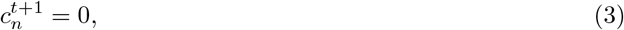

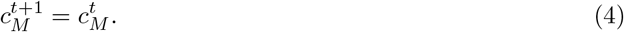

where *c*^*t*^ *∈ C*^*t*^ denotes the clone *c* at time step *t*. The next state of the simulation is then obtained as *C*^*t*+1^ = {*c*^*t*+1^ | *c*^*t*^ *∈ C*^*t*^}.

Note that if *p*_birth_ > *p*_death_ the population size would increase exponentially. In the opposite case, the population would eventually die out. In this homeostatic model the birth and death probability are equal to the turnover probability, therefore the population size remains constant on average. However, due to stochastic fluctuations, the population will always die out after a large, but finite number of steps (extinction event). This is common to all frameworks that include stochastic cell death (Roch, 2015) and the extinction probability grows with *θ*_turn_ (Supp. Fig. 1). We avoid the problem in practice by setting a sufficiently small *θ*_turn_ and a correspondingly large *min pop*.

In summary, at this stage, our system comprises a single clone, which is roughly constant in size and shows no clonal dynamics.

### 2.3 Mutations

We next introduce mutations into the homeostatic model. We assume that each mutation is unique (infinite sites assumption) and therefore each mutation spawns a new clone. New mutations can only occur during the division step, with the probability *θ*_mut_ per daughter cell (2*θ*_mut_ per cell division). For simplicity we limit the number of mutations to at most half the daughter cells, so that the size of existing clones does not decrease due to cell division. As we only focus on driver mutations in biologically realistic scenarios, we expect *θ*_mut_ ≪ 1 and thus this limitation does not affect the simulation in practice. We then extend (2) to:

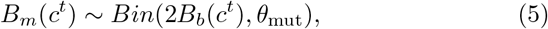

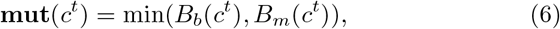

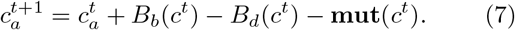

Mutated cells are removed from the group of new cells (7) and each of them spawns a new clone each with exactly one alive cell, such that for each *c*^*t*^ *∈ C*^*t*^:

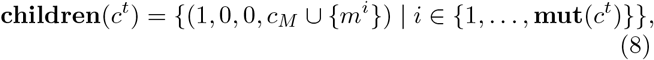

where *m*^*i*^ is a new, unique mutation. The new clones are then added to the updated population, i.e. *C*^*t*+1^ = {{*c*^*t*+1^} ∪ **children**(*c*^*t*^) | *c*^*t*^ ∈ *C*^*t*^}.

Our homeostatic system now consists of several clones, none of which has a fitness advantage over the over. In such a system the appearance and disappearance of clones is simply due to stochastic fluctuations and constant accumulation of new neutral mutations (Fig. 1).

**Figure 1:**
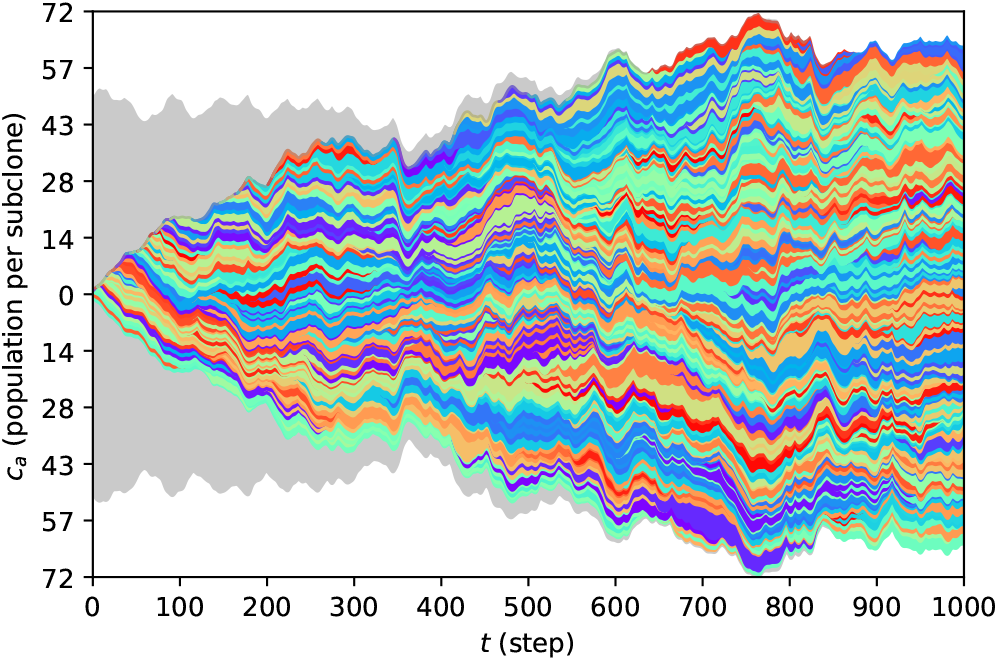
An illustration of mutation accumulation over 1000 steps with homeostatic population of 100 cells (*C*^0^ = {(100, 0, 0, {*m*})}) and the mutation probability *p*_*mut*_ = 0.25. Each colour represents the size of a population of a single clone. We can see that the original population (grey) disappears as a result of mutation accumulation during turnover. Some of the newly created clones disappear as well while others branch into further clones.

### 2.4 Fitness and model variants

We now abandon the homeostatic scenario by introducing fitness. Traditionally, mutations are considered to be either fitness-increasing *drivers* or neutral or almost-neutral *passengers*. Driver mutations increase the evolutionary advantage of a cell (Alberts *et al*., 2002), represented by a positive change in cellular fitness (Fu *et al*., 2022; Noble *et al*., 2022; Beerenwinkel *et al*., 2015). Here we focus on how the growth behaviour changes with fitness increasing mutations and omit passenger mutations.

The fitness increase *F* (*m*) that a new mutation *m* provides is drawn from a fitness distribution controlled by the model option *o*_dis_ ∈ {*uni, exp, norm, const*}, referring to the uniform, exponential, truncated normal, and constant distributions, respectively. We here use the term constant distribution as a shorthand for a single-element discrete uniform distribution. All possible fitness distributions are defined such that their with mean at *θ*_fit_:

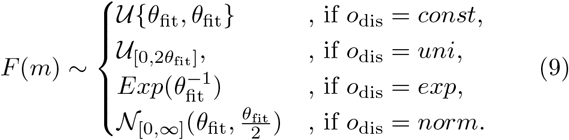

Here, 𝒩 _[0,*∞*]_ represents the truncated normal distribution for which we set the lower bound at 0. For standard deviation we use 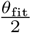 in line with Bozic *et al*. (2010). Note that for a lower bound of 0 and 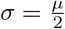 the expected value is 1.027 *·θ*_fit_. The exact shapes of the individual distributions are shown in Supp. Fig. 2.

Since a clone inherits the mutations of its parent clone, the fitness values of multiple mutations have to be combined through the use of an accumulation function controlled via the accumulation option *o*_acc_. Three accumulation functions are considered: multiplicative (*mul*), limited multiplicative (*lim*), and additive (*add*). The limited multiplicative accumulation function has been implemented according to Noble *et al*. (2022) and its full formulation is in Supp. Fig. 3. We obtain the joint fitness for all the mutations in mutations 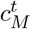 using the accumulation function

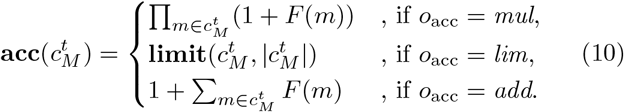

Lastly, we consider three options of applying the fitness *effect* to a clone: fitness increases the birth probability (*birth*); fitness decreases the death probability (*death*); or the fitness is split equally between the birth and death probability (*both*), as also used in Watson *et al*. (2020). We then calculate the **birth** 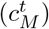 and **death** 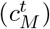 effect functions as follows:

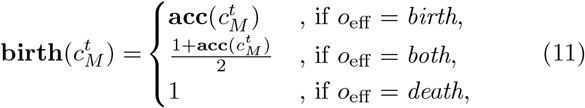

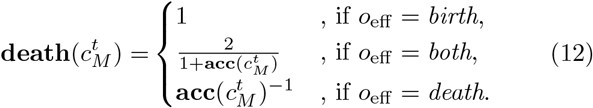

The birth and death probabilities are then given as:

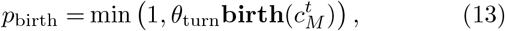

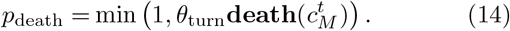

The homeostatic scenario is a special case of the above, since if *θ*_fit_ = 0 then *p*_birth_ = *p*_death_ = *θ*_turn_. Unless stated otherwise, we use *o*_dis_ = *const* and *o*_acc_ = *mul* as default values (Table 1).

With the fitness change now included our system describes a well-mixed population of cells without spatial constraints.

### 2.5 Confinement

We now introduce quasi-spatial constraints into our model of a well-mixed population of cells.

Driver mutations inevitably lead to exponential growth in the absence of limiting factors. In cancer, these factors include e.g. lack of access to blood vessels and nutrient limitations or spatial constraints (Folkman, 1971). We represent these constraints in an abstract manner using the *confinement* parameter *θ*_conf_, which limits the turnover of cells by the size of the tumour (Fig 2). Confinement acts in two ways. First, it limits the number of cells that can divide based on the size of the tumour. Second, it prevents some cells from disappearing after cell death, instead turning them into necrotic cells instead which continue to contribute to the size of the tumour.

**Figure 2:**
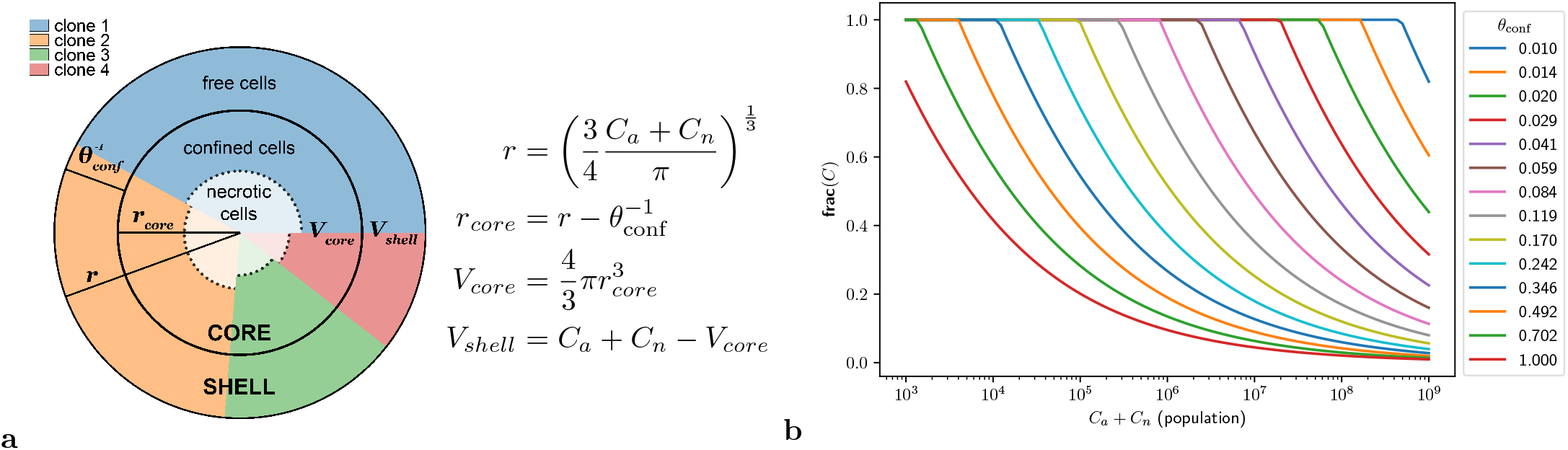
Confinement: **a)** Confinement splits the tumour into a proliferating shell (free cells) and a core with radius *r*_*core*_, composed of confined and necrotic cells. Different colours represent populations of clones. Each cell is expected to take a unit volume in the densely packet tumour. The probability of a cell diving is then a combination between its fitness and the probability of it being in the shell. Similarly, the dead cells are either removed or become necrotic based on the shell fraction. **b**) Illustration of the **frac** function across the range of the confinement values and population sizes. For this plot we set *C*_*a*_ = *C*_*n*_.

To formulate the confinement, we create a geometrical representation of the tumour as a sphere (Fig 2a). We fix the spatial scale of our model such that each individual cell has unit volume and the volume of the whole tumour at the time *t* is equal to the number of its alive and necrotic cells 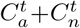. The tumour is thus divided into two regions, a proliferating *shell* and a quiescent *core*. Cells in the shell can divide and get removed when they die. Conversely, cells in the core cannot divide due to lack of space or resources, and turn into necrotic cells upon death. We denote **shell-V**(*C*^*t*^) the volume of the shell and **core-r**(*C*^*t*^) the radius of the core for the population contained in *C*^*t*^. Under the assumption of a perfect sphere, we can compute the fraction **frac**(*C*^*t*^) of the tumour volume occupied by the shell, relative to the shell volume **shell-V**(*C*^*t*^) and the radius of the core **core-r**(*C*^*t*^) as follows:

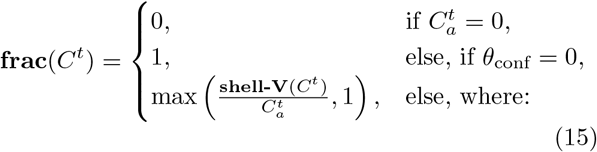

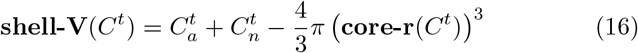

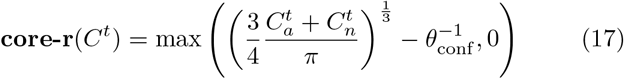

Note that the width of the shell is given by 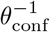, meaning that lower confinement values lead to a larger shell and subsequently a larger proportion of proliferating cells (Fig 2b). In particular, for *θ*_conf_ → 1 we approximate the boundary of the sphere, i.e. its surface, while for *θ*_conf_ → 0 the whole sphere is considered the shell, irrespective of its size. We can therefore easily emulate surface growth conditions (*θ*_conf_ = 1), volume growth conditions (*θ*_conf_ = 0), or any mixture of the two, without explicitly tracking the position of individual cells. Combining the above we then obtain the final model:

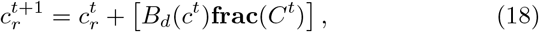

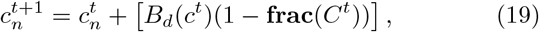

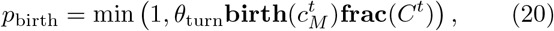

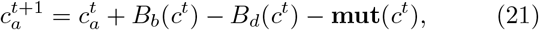

where [·] refers to rounding to the nearest integer. While for the birth probability (20) directly depends on the fraction (15) to ascertain that even newly spawned single-cell clones can divide, the death probability (14) does not depend on the fraction. There the fraction only separates removed from necrotic cells.

As only the *o*_acc_ = *lim* option has an upper boundary, a situation where **birth** 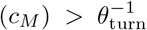 ^1^ is possible. Consequently, due to the upper boundary *p*_*birth*_ ≤ 1, the confinement may play a role in establishing the upper limit for fitness. However, for the default values *θ*_*fit*_ = 0.2, *θ*_turn_ = 0.01, this only becomes a factor when |*c*_*M*_ | > 25, while even in most extreme cases we do not expect more than 20 driver mutations for biologically plausible parameter values.

Note that if we set *θ*_conf_ = 0, the model is equal to the one in Sec. 2.4, with also *θ*_fit_ = 0 to Sec. 2.3, and with *θ*_mut_ = 0 to Sec. 2.2.

### 2.6 Parameter space

We aim to simulate realistically-sized tumours with the population size ∼10^9^ cells, corresponding to a tumour of > 1*cm*^3^ in size (Alberts *et al*., 2002; Del Monte, 2009). We set the minimum population size *min pop* for a simulation to be considered to 1000 cells. The *max_pop* is then derived from *min_pop* as 1000 · 2^20^ ∼ 10^9^, hence population doubling occurs 20 times starting from *min_pop*.

To calibrate overall cellular turnover, we created a homeostatic model without mutations and a starting population size of 100 cells and evaluated the turnover probability in the following steps: *θ*_turn_ ∈ {0.1, 0.05, 0.01, 0.005} (Supp. Fig. 1). We found that for *θ*_turn_ = 0.01 there was no extinction event within *max steps* = 1000 for any of the 1000 replicates and we thus selected this as a turnover value for all further simulations to balance granularity of the simulation and execution time.

We fixed the limiting variable for the maximum number of generations *max steps* to 10^6^, such that without a mutation the initial cell divides on average 10^6^ *·* 0.01 = 10^4^ times. For a proliferating cell where typical division cycle is 24h (Alberts *et al*., 2002), this would correspond to ∼ 27 years of real time. In all subsequent simulations the execution reached the maximum population size before reaching the maximum number of generations.

We explore the choice of values for the confinement *θ*_conf_, mutation probability *θ*_mut_ and average fitness increase *θ*_fit_ parameters in the Results section (Section 3.1).

### 2.7 Population metrics

To evaluate the individual simulation runs and to compare their clonal behaviour to prior work as well as experimental data, we implement three metrics: the *mean number of drivers per cell*, the *clonal diversity index*, and the *clonal fluctuation score* which characterises clonal diversity over time.

To speed up computation of summary statistics we only consider clones larger than a minimum fraction *cutoff* of the population 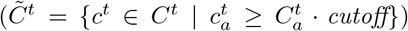 and only calculate the metrics at every time step at which the population of alive cells doubles. The Fish plots (e.g. Fig. 3 a-c) are not affected by the cutoff, but only display clones that reached at least 1% of the population at some point.

**Figure 3:**
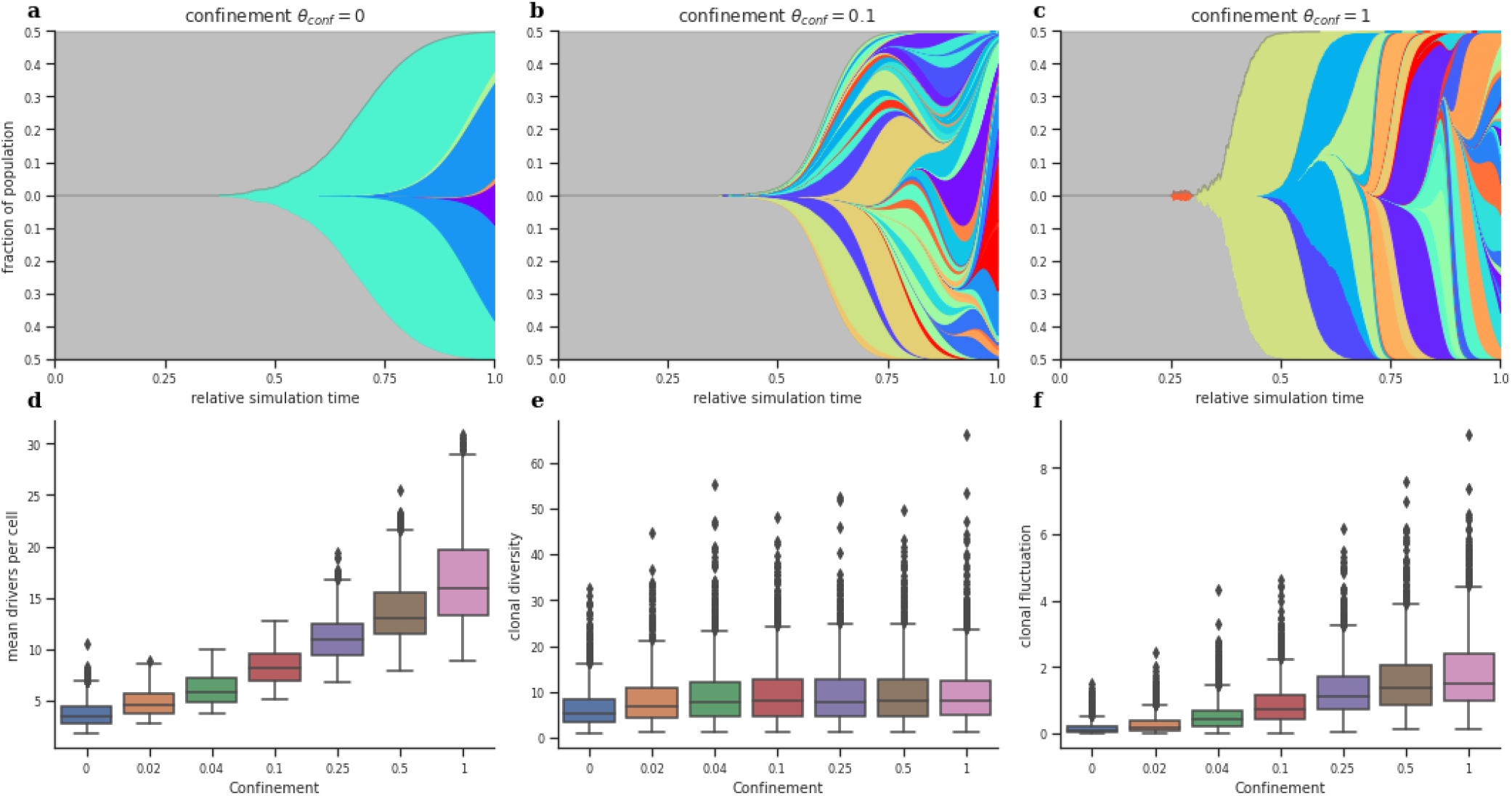
Simulations with confinement:**a-c)** Selected fish plots for minimum, intermediate, and maximum confinement values. **a)** In the absence of confinement (*θ*_conf_ = 0), corresponding to well-mixed volume growth, we find a limited amount of clonal diversity and few drivers. **b)** For intermediate confinement (*θ*_conf_ = 0.1) we find an increase in the number of co-existing clones as well as an increase in the number of drivers. **c)** For the maximum confinement (*θ*_conf_ = 1) we observe increase in clonal sweeps, increasing the number of mutations, but not the diversity. **d-f)** Increasing the confinement continuously increases the final number of mean drivers per cell and the clonal fluctuation, while the clonal diversity plateaus around *θ*_conf_ = 0.1. For every confinement value the data is aggregated over all values of the mutation probability and the mean fitness increase.

The *mean number of drivers per cell* 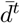 tracks the mutational burden of the growing tumour and is defined as

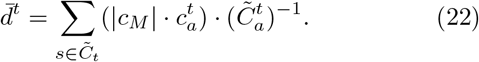

The *clonal diversity index D*^*t*^ reflects the total number of clones and their size and is based on the inverse Simpson index defined as

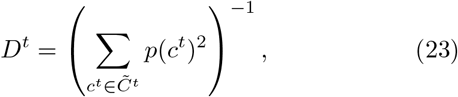

where 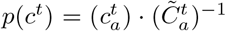 is the fraction of the total population of clone *i* at time step *t*. As shown in Noble *et al*. (2022), this measure has a lower boundary of 1 and is robust to the presence or absence of small populations. If there are *k* clones with equal population sizes, then *p*(*c*^*t*^) = *k*^−1^ for all 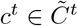 and 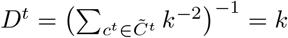.

Since *D*^*t*^ is a static measure of clonal diversity at time point *t*, we also consider the *clonal fluctuation score D*_fluc_ as the difference between the mean absolute slope minus the mean slope of the clonal diversity over time:

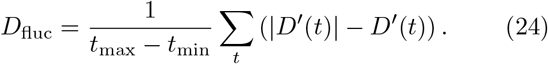

If the clonal diversity is monotonously increasing, then *D*_fluc_ = 0.

Estimates for the population metrics can be retrieved from Noble *et al*. (2022) who calculated metrics for single-cell experiments of spatially organised tumours to be in the following ranges: clonal diversity *D*^*t*^ ∈ [1, 12] and mean number of drivers per cell 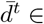. The latter was confirmed with a measured values of 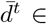] (Bozic *et al*., 2010). Unfortunately, due to the lack of longitudinal data for cancer, there are no current estimates for the clonal fluctuation score.

### 2.8 SMITH implementation

SMITH has been implemented in C# as an open-source package under the MIT license and is available at bitbucket.org/schwarzlab/smith with pre-compiled binaries for Windows, Linux, and MacOS. The exact code used for producing the results and figures in this manuscript is available at doi.org/10.5281/zenodo.6885041. The Fish (Muller) plots were implemented in the accompanying open-source Python library *PyFish*, available at bitbucket.org/schwarzlab/pyfish, through PIP, and on Bioconda.

Using SMITH, simulating a population with 10^9^ cells took approximately a minute on a single-threaded CPU using less than 1GB of memory.

## 3 Results

### 3.1 Confinement replicates dynamics of spatial models

Previous research has shown that non-spatial models produce tumours with a low number of drivers (between 1–4), and limited clonal diversity of 3 or fewer clones of roughly equal size (Noble *et al*., 2022), in line with observations in non-spatial tumours such as acute myeloid leukaemia (AML). In contrast, spatially organised tumours generally demonstrate far greater clonal diversity (*D*^*t*^ ∈ [1, 12]) and a larger number of drivers 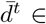, an observation that could so far only be recreated by explicitly spatial models of tumour evolution (Noble *et al*., 2022).

We hypothesised that our confinement mechanism alone should be sufficient to reproduce the clonal diversity and high number of drivers observed in spatially organised tumours. To test this, we first conducted a grid search of varying values of the confinement parameter *θ*_conf_, the mutation probability *θ*_mut_ and the average fitness increase *θ*_fit_. Each of the parameters was varied over the log-scaled range of *θ*_conf_ ∈ [0, 1], *θ*_mut_ ∈ [2.5 ·10^−5^, 4 ·10^−4^] and *θ*_fit_ ∈ [0.05, 0.8]. Driver mutations were set to a constant fitness increase (*o*_dist_ = *const*), acting on the birth probability (*o*_eff_ = *birth*), and fitness values of multiple mutations were accumulated using multiplication (*o*_acc_ = *mul*). We conducted 50 simulations for each value combination and inspected the clonal architectures with respect to the three evolutionary metrics: mean number of driver mutations, clonal diversity index, and clonal fluctuation score (Methods and Supp. Fig. 5,6). For visualisation and analysis, we use the marginal effects of each parameter across the full range of all other parameters. For the precise values for each combination please refer to Supp. Fig. 5.

In the absence of confinement (*θ*_conf_ = 0), all cells in the tumour can divide and we observed a behaviour reminiscent of well-mixed non-spatial cell populations (Fig. 3a, Supp. Fig. 4a). Overall genetic diversity was low (median clonal diversity ∼ 5.4, 25-75 quantiles: 3.3-8.4), only few driver mutations occurred (median of 3.5, 25-75 quantiles: 2.8-4.5), and virtually no clonal fluctuations were visible (median of 0.1, 25-75 quantiles: 0.0-0.2) (Fig. 3d-f). Due to the fitness increasing effect of mutations on the birth rate, the early appearing clones remained in the population until the end of the simulation was reached at 10^9^ cells. Note that the observed value for the diversity is slightly higher than observed in similar non-spatial models (Noble *et al*., 2022) which is explained by the choice of our fitness function and elaborated in Sec 2.4.

Increasing the confinement parameter to *θ*_conf_ = 0.1 limits cell division to the outer 10 cell layers (Methods) in a spherical tumour model (Fig. 3b, Supp. Fig. 4b). As a result we observed an overall increase in the mean number of drivers (median 8.1, 25-75 quantiles: 6.9-9.6) and a substantially higher clonal diversity (median 8.0, 25-75 quantiles: 4.8-12.6), in keeping with the literature about observations in spatially organised tumours (Noble *et al*., 2022). We also observed an increase in the clonal fluctuation score (median of 0.7, 25-75 quantiles: 0.4-1.2) (Fig. 3d-f). We argue that this increase in both driver accumulation and diversity is due to an increased evolutionary pressure caused by competition for space, imitated by our confinement mechanism, which slows the growth of existing clones and allows for new clones to appear and founder clones to disappear in favour of late, high-fitness clones.

Using the maximum confinement (*θ*_conf_ = 1) limits cell division to the outermost layer of a spherical tumour (Methods, Fig. 3e, Supp. Fig. 4c). As a result, we observed a stark increase in the mean number of drivers (median 15.9, 25-75 quantiles: 13.3-19.7) whereas the clonal diversity remains similar at a median value of 8.0 (25-75 quantiles: 4.9-12.5). We also observe a further increase in the clonal fluctuation score (median of 1.5, 25-75 quantiles: 1.0-2.3) (Fig. 3d-f). Across the full parameter range, confinement leads to an approximately logarithmic increase in the number of drivers (Fig. 3d). While initially, this also leads to an increase in clonal diversity (Fig. 3e), the increase in evolutionary pressure by higher confinement values also increases the likelihood of clonal sweeps, where a clone quickly outcompetes its neighbours and captures the whole population, reducing clonal diversity once again. This effect is clearly visible in the increase of the clonal fluctuation score (Fig. 3f), but leads to clonal diversity remaining static beyond moderate values of confinement (Fig. 3e).

As expected, we find that increasing the mutation probability *θ*_mut_ simply accelerates clonal evolution, leading to a logarithmic increase in all our analytical metrics (Supp. Fig. 6-left column). As all clones grow exponentially, increasing the rate at which new mutations appear allows the new clones to faster reach the same population as clones from the previous generation, increasing the rate of accumulation of new drivers and likelihood of clonal sweeps. In contrast to the mutation probability, we find that increasing the average fitness advantage of a driver mutation (*θ*_fit_ ∈ [0.05, 0.8]) decreases the the clonal diversity and clonal fluctuation scores (Supp. Fig. 6e,h). The mean number of drivers is only weakly affected by changes to the average fitness advantage with median values all in the range of 7.7-8.9 (Supp. Fig. 6b). For a value of *θ*_fit_ = 0.05 gaining an additional driver increases the fitness of the clone by a factor of 1.05 (Eq. 10). The newly created clone therefore possesses fitness similar to its predecessor, allowing for co-existence and the observed high clonal diversity. For the much higher value of *θ*_fit_ = 0.8, gaining an additional driver almost doubles the fitness of the clone (factor of 1.8), which then rapidly outgrows the existing population, leading to a less diverse tumour with a ladder-like evolutionary tree. This is supported further by the behaviour of the mean drivers per cell with varying average fitness values (Supp. Fig. 6b), as the median value of the mean drivers per cell remains unchanged but the standard deviation falls from 7.1 for *θ*_fit_ = 0.05 to 2.7 for *θ*_fit_ = 0.8

By comparing the observed test metrics to ones calculated for real tumours as shown in Sec. 2.7, we choose the final parameter set *θ*_mut_ = 10^−4^, *θ*_fit_ = 0.4 and *θ*_conf_ = 0.1. The selected *θ*_mut_ has been used in other publications: *θ*_mut_ ∈ [10^−3^, 10^−4^] in (Fu *et al*., 2022) and *θ*_mut_ ∈ {10^−4^, 10^−5^, 10^−6^} in (Noble *et al*., 2022). The chosen confinement value of 0.1 leads to a mixture of surface growth and volume growth, where the ten topmost layers of the spherical tumour can divide. For an overview of the final parameters and variables see Table 1.

In summary, in the absence of any confinement SMITH recreates previous findings of well-mixed models that resemble non-spatial tumours such as leukemia. In contrast, out results show that confinement alone is sufficient to create the rich clonal dynamics otherwise only seen in explicitly spatial simulations and in spatially organised tumours.

### 3.2 Clonal dynamics are complex and variable over time

Most simulation studies so far have focused on interpreting the simulation endpoint after a certain number of generations and have largely avoided investigations into the variability of repeated simulations conducted with the same set of parameters. To address this and to learn about the progression of the clonal dynamics over time (Fig. 4a-c, Supp. Fig. 7), we queried repeated simulation runs at fixed time points indicated by each doubling of the population size and investigated the trajectories of our metrics over time, motivated by the observation that across cancer types the doubling time remains roughly constant (Talkington and Durrett, 2015).

**Figure 4:**
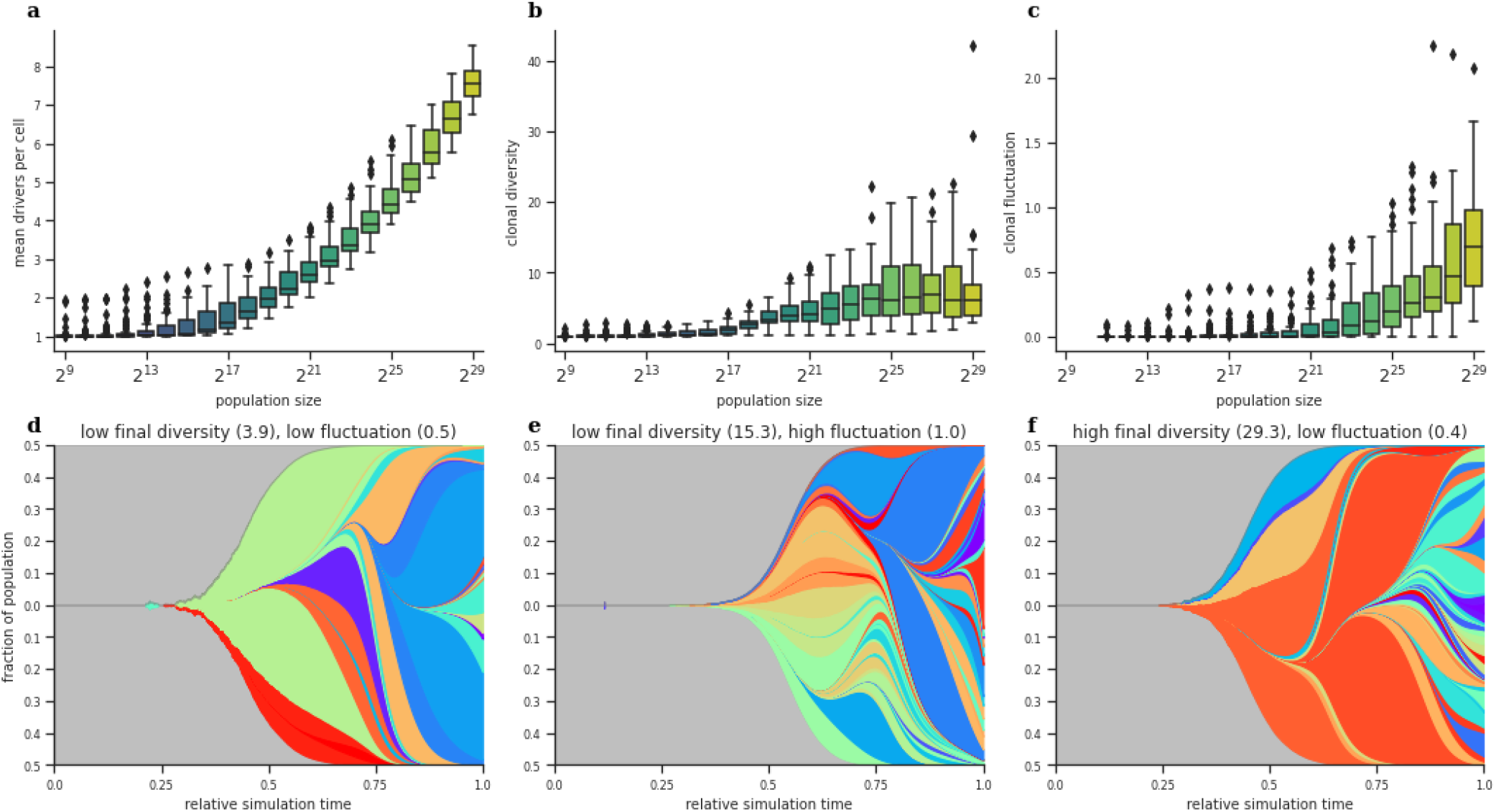
Population metrics over time (parameters according to Table 1):
**a-c)** Both the mean number of drivers per cell and the clonal fluctuation increase continuously with the growing population size throughout the simulation. In contrast, the clonal diversity plateaus for large population sizes. **d-f)** Selected Fish plots for simulations with the same parameter set. We see clear heterogeneity in the population dynamic despite the limited amount of stochasticity. **d)** This example exhibits low clonal diversity throughout its evolution. **e)** Here we observe a burst in diversity halfway through the simulation followed by a clonal sweep of the blue clone. **f)** In this final example, the simulation is dominated by a few clones until the final fifth of the simulation, where a burst of new clones leads to a high final diversity.

We observed that the mean drivers per cell grows steadily with increasing population size (Fig. 4a), demonstrating that newly appearing clones eventually overtake the population. Clonal diversity initially surges until the population size of ∼ 2^20^(∼ 10^6^) cells is reached, after which the increase decelerates and finally reaches a plateau around 2^27^(∼ 10^8^) cells (Fig. 4b). Below 2^16^ cells there is on average less than 1.5 drivers per cell, as the first clone is dominant in the population. Only when the tumour is large enough for new clones to appear and once these clones start competing with the first clone do we also see an increase in diversity.

Conversely, we observe a continuous increase in the clonal fluctuation score with increasing size of the tumour (Fig. 4c), suggesting continued clonal dynamics throughout the lifetime of a tumour, even when the clonal diversity has reached its maximum. To investigate the dynamics of the clonal fluctuation score in more detail, we looked at individual trajectories of the clonal diversity score over time for different simulation runs with the same set of parameters. We found that the trajectories leading to the final values are very diverse (Supp. Fig. 8) and we observed several distinct patterns. For example, some simulated tumours maintained an overall low diversity due to repeated clonal sweeps and a correspondingly low fluctuation score overall (Fig. 4d). Other tumours demonstrated a continuous rise and fall in diversity, where clonal expansions and clonal sweeps acted interchangeably, leading to an overall high fluctuation score (Fig. 4e) and medium average clonal diversity. On the other end of the spectrum we observed a monotonous increase in the number of coexisting clones, leading to a high clonal diversity with simultaneously low clonal fluctuations (Fig. 4f).

Interestingly, clonal sweeps, which are an important characteristic of tumour evolution (Black and McGranahan, 2021), and which naturally emerge in our confinement-based model, have not previously been described even in explicitly spatial simulations (Noble *et al*., 2022; Fu *et al*., 2022; Chkhaidze *et al*., 2019; Sottoriva *et al*., 2015).

In summary, while we observe a common trend towards higher number of drivers and plateauing clonal diversity with increasing size of the tumour, we also observe extensive heterogeneity in the trajectories of clonal diversity and fluctuations over time, governed by alternating periods of clonal sweeps and diversification.

### 3.3 The effect of the fitness function on clonal evolution

Prior work has claimed that the choice of the fitness function does not have significant bearing on the outcome of the simulation (McFarland *et al*., 2013; Chkhaidze *et al*., 2019; West *et al*., 2021). To test this claim, we implemented four fitness distributions (constant, uniform, normal, and exponential) with the same mean fitness advantage and compared them with respect to our clonal dynamics metrics (Methods).

We found that the choice of fitness distribution substantially affects the outcome of our simulation (Fig. 5). For example, the constant fitness gain used above yields the maximum number of mean drivers per cell (median 7.3, 25-75 quantiles: 7.0-7.7, Fig. 5a) as well as the highest amount of clonal diversity (median 9,3, 25-75 quantiles: 5.6-15.1, Fig. 5b) and clonal fluctuation (median 0.3, 25-75 quantiles: 0.2-0.4, Fig. 5c). In contrast, normal and uniform distributions showed substantially lower values compared to the constant fitness function (median 6.5 and 6.7 mean drivers per cell, median 5.3 and 4.8 clonal diversity and median 0.2 and 0.3 clonal fluctuation for normal and constant distributions respectively), while the exponential distribution distinctly exhibited the lowest number of clonal diversity (median 3.0), fluctuation (median 4.9) and number of drivers (median 0.5).

**Figure 5:**
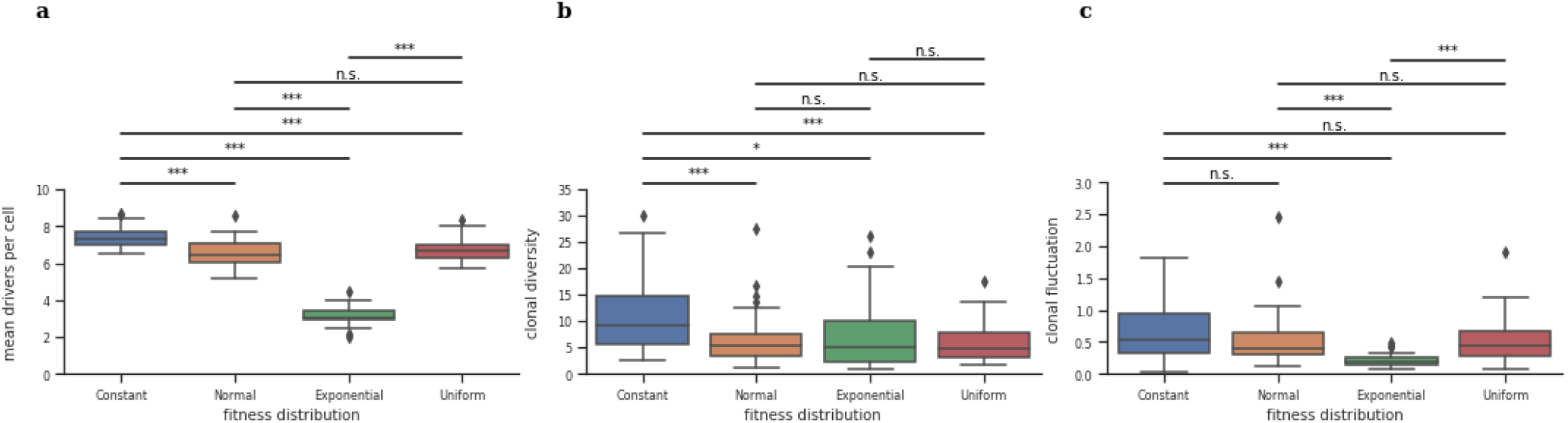
Effect of the model fitness options:**a)** Except for the normal and the uniform distribution, all distribution options lead to a differing range of mean drivers per cell with the constant function leading to the highest number of drivers. **b)** The constant function leads to a significantly higher clonal diversity compared to the other distributions. **c)** The exponential distribution leads to a significantly lower clonal fluctuation compared to the other distributions. *: *p <* 0.05, **: *p <* 0.005, ***: *p <* 0.0005, n.s.: not significant.

The difference between the constant fitness gain and the three other fitness distributions can be explained by the fact that two clones with the same number of drivers have the same fitness and therefore do not have an evolutionary advantage over each other, allowing for coexistence. In contrast, using non-constant fitness functions invariably leads to expansions of a small number of clones with a comparatively high fitness advantage. In particular, when using the longtailed exponential distribution, a vastly more potent mutation will occasionally appear, sweeping through the population. As a side effect, the simulation with an exponential function is considerably shorter as these super-clones divide faster than those with mean fitness values (Supp. Fig. 9j).

In addition to the fitness distribution itself, the fitness increase per generation is modulated by the accumulation function used to aggregate the fitness values of multiple mutations, and whether a fitness change affects the birth rate or death rate of cells or both. We tested the effects of three different accumulation functions (additive, multiplicative, and multiplicative with an upper bound) on the cellular birth rate, death rate, or both (Sec. 2.4).

We find that, in contrast to the choice of the fitness distribution, the choice of accumulation function and its effect on cell birth or death does not have a significant impact on the clonal diversity (Supp. Fig. 9e,f). However, we did observe a difference in the amount of drivers per cell (Supp. Fig. 9b), where the multiplicative accumulation function generates the highest number of drivers (median 7.3, quantiles: 7.0-7.7) followed by the multiplication with an upper bound (median 6.9, quantiles: 6.7-7.2) and finally the additive accumulation function (median 6.0, quantiles: 5.8- 6.2). Multiplicative accumulation leads to a higher fitness for the same number of drivers compared to the other two accumulation methods, which results in a higher turnover rate and therefore a higher chance to acquire new drivers.

We also found changes in fitness that affect the cell birth probability to lead to the highest number of mean drivers per cell (median 7.3, 25-75 quantiles: 7.0-7.7), followed by fitness changes that affect both birth and death probability (median 6.6, 25-75 quantiles: 6.4-6.8) (Supp. Fig. 9c). Fitness changes that affect only the death probability lead to the lowest number of mean drivers per cell (median 5.5, 25-75 quantiles: 5.4-5.6). This choice also has a weak effect on the clonal fluctuation score (Supp. Fig. 9h,i). The most aggressive variants (multiplicative, birth) seem to exhibit the lowest fluctuation. This can be explained by the fact that the most aggressive variants reach the target population size in considerably fewer steps (Supp. Fig. 9k,l), and have therefore less time to fluctuate. In this sense the clonal fluctuation seems to be quite robust with respect to the overall number of simulation steps and with lower turnover parameter the difference would likely become insignificant.

In summary, We find that the choice of fitness distribution, but not the choice of accumulation function or where the fitness change is applied (birth, death or both), widely affects the clonal dynamics of the simulated tumour mass, in contrast to existing reports, and should be chosen carefully when making modelling decisions.

## 4 Discussion

We have presented SMITH, a stochastic model of tumour evolution which implements confinement, a novel mechanism adding spatial constraints to the non-spatial branching process model of cancer. Confinement induces evolutionary pressures through competition for space by splitting the tumour into a proliferating shell and a static inner core. It thereby generates clonal dynamics traditionally only observed in explicitly spatial models and spatially organised tumours. While splitting a tumour into a replicating and non-replicating part has been previously used in algebraic descriptions of tumour cell population size (Paterson *et al*., 2016; Dassios *et al*., 2012), to the best of our knowledge SMITH is first to integrate such a principle into an evolutionary model.

Naturally, multiple different formulations of the confinement mechanism are conceivable. The one presented here has the advantage of allowing for a direct geometric interpretation in terms of tumour shell volume under the assumption of a tumour as a perfect sphere. While not all real-world tumours are spheroids, other formulations based on different shapes are easily implementable, e.g. to accommodate discoidal or segmental shapes (Byrd *et al*., 2020). In particular, in Talkington and Durrett (2015) the authors claim a growth rate with an exponent of 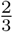 as typical for solid tumours. The exponent of 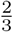 also (when multiplied by an appropriate constant) projects a volume of a 3D object onto its surface, and really any function that projects a 3D object onto its surface area could be considered.

A major advantage of the confinement approach is computational speed, which is several orders of magnitude faster than explicitly spatial models of tumour growth. Such explicitly spatial models are commonly run on high-performance computers in parallel, with cell populations in millions of cells or lower (Rosenbauer *et al*., 2020). In contrast SMITH performs a single simulation up to 10^9^ cells in minutes on a desktop computer, enabling detailed analysis of the variability of evolutionary trajectories under a variety of different parameterisations. This efficiency naturally comes at the expense of complexity and as such SMITH currently does not capture some features of real-world tumours, such as cell migration or interactions with the immune system or tumour microenvironment.

We purposefully kept the stochasticity of our model low and limited to fluctuations in population size and the timing of new mutations. Despite this limited stochasticity, we observed for a single set of parameters a wide variety of heterogeneous evolutionary trajectories, where tumours would switch between periods of relative stasis and clonal diversifications in a seemingly erratic manner. Based on these results we argue that predictions of the future of an evolving tumour based on its past and present, an important topic for current cancer evolution research, might prove to be more challenging than anticipated.

When building models, specific modelling choices are ideally based on real-world data. While our estimates for cell turnover and mutation rates are well in keeping with the literature, there is no clear consensus on how fitness changes are distributed, how fitness values should be accumulated across multiple mutations, and how these choices affect the birth and death rate of cells. We tried to explore these choices in concert and, in contrast to previous findings (McFarland *et al*., 2013), we observed that the shape of the fitness distribution has a substantial impact on the results of the simulation, in particular when comparing long-tailed and short-tailed distributions. Additional research in these areas will be needed and ought to be complemented by experimental data.

In summary, we have shown that our simple, high-performance simulation model provides an effective way of observing the fundamentals of cell-based exponential growth with fitness-increasing mutations and spatial constraints. SMITH thereby produces results in close agreement with observations on real-world tumours including recapitulation of specific clonal dynamics such as interchanging periods of clonal sweeps and diversification previously unseen in non-spatial models of tumour evolution. In future work, more complex variations of this model are conceivable by adding biologically relevant mechanisms such as interactions with non-malignant cell populations and cell exchanges between multiple tumour masses to mimic metastatic seeding.

## Acknowledgements

The authors would like to thank Klaus-Robert Müller for support. Computation has been performed on the HPC for Research cluster of the Berlin Institute of Health.

## Author Contributions

AS and TK conceptualised the method. AS, TK and RFS designed the model. AS and TK implemented the model. AS, TK and RFS wrote the manuscript. RFS supervised the project.

## Funding

This work was partially funded by the German Ministry for Education and Research as BIFOLD - Berlin Institute for the Foundations of Learning and Data (ref. 01IS18025A and ref 01IS18037A). RFS is a Professor at the Cancer Research Center Cologne Essen (CCCE) funded by the Ministry of Culture and Science of the State of North Rhine-Westphalia. The authors would like to thank the Helmholtz Association (Germany) for support.

## Notes

### Competing Interest Statement

The authors have declared no competing interest.

