## Supplementary Material for "SMITH: Spatially Constrained Stochastic Model for Simulation of Intra-Tumour Heterogeneity"


Adam Streck[1], Tom Kaufmann[2,1], and Roland F. Schwarz[3,2,1]

[1] *Berlin Institute for Medical Systems Biology, Max Delbrück Center for Molecular Medicine in the Helmholtz Association, Berlin, Germany*

[2] *BIFOLD - Berlin Institute for the Foundations of Learning and Data, Berlin, Germany*

[3] *Center for Integrated Oncology, Cancer Research Center Cologne Essen, Faculty of Medicine and University Hospital Cologne, University of Cologne, Cologne, Germany*

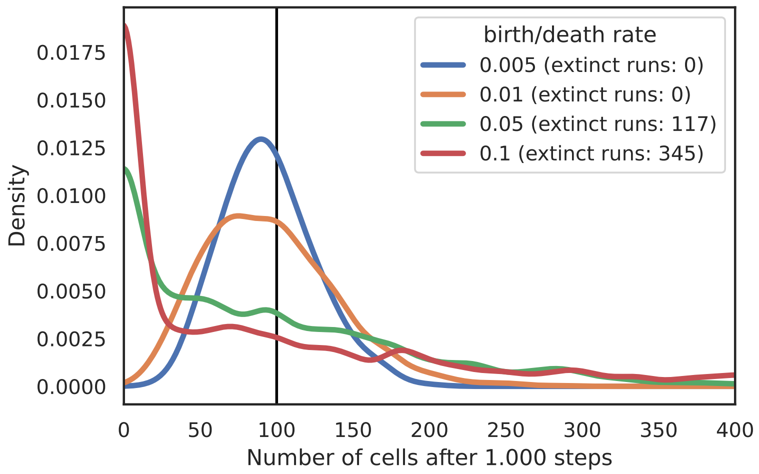


Figure 1: Homeostasis without mutations: We simulate 1000 runs each for four separate values of $\theta_{turn}$ each starting at 100 cells. For 0.05 and 0.1 the turnover is high enough that we observe extinction of the population in multiple runs.

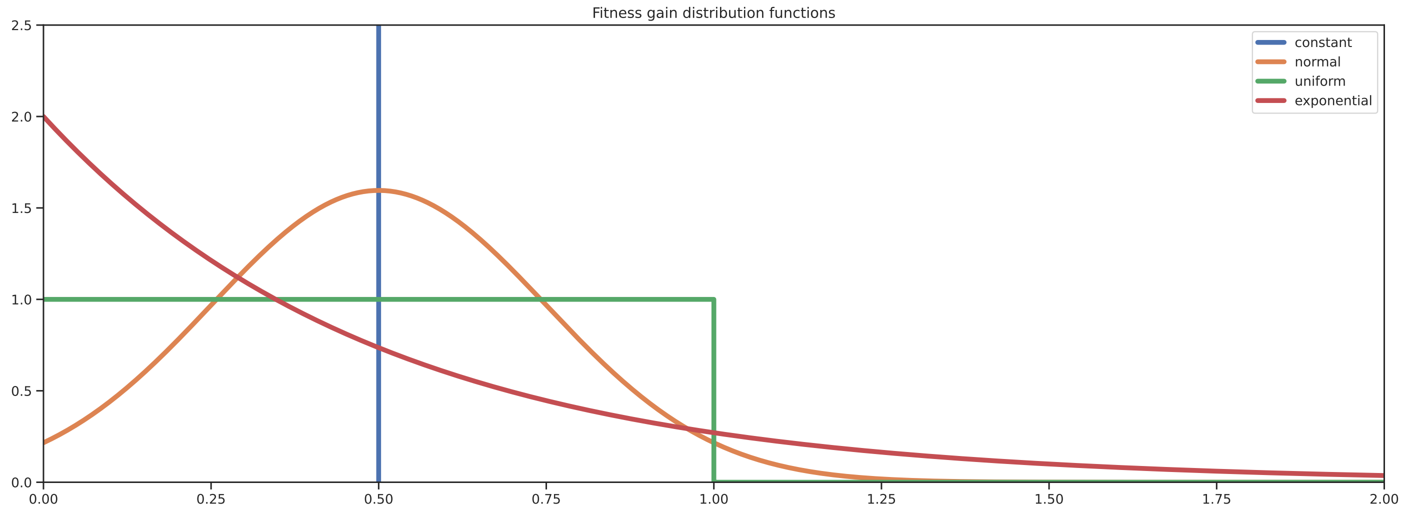


Figure 2: Distribution of fitness around the mean $\theta_{fit} = 0.5$ for the random variables $C^{\theta}$ (blue), $N^{\theta}$ (orange), $U^{\theta}$ (green), and $E^{\theta}$ (red).

$$\mathbf{limit}(c_M, i) = \begin{cases} 1 & \text{, if } i \leq 0, \\ \mathbf{limit}(c_M, i-1)\left(1 + F(c_{M(i)})(1 - \frac{\mathbf{limit}(c_M, i-1)}{10})\right) & \text{, else.} \end{cases}$$

Figure 3: The limited multiplicate function **limit** given in (Noble *et al.*, 2022) The function is calculated dependent on the order in which mutations have been accumulated. To this end we use the notation $c_{M(i)}$ to denote the $i$-th mutation in $c_M$.

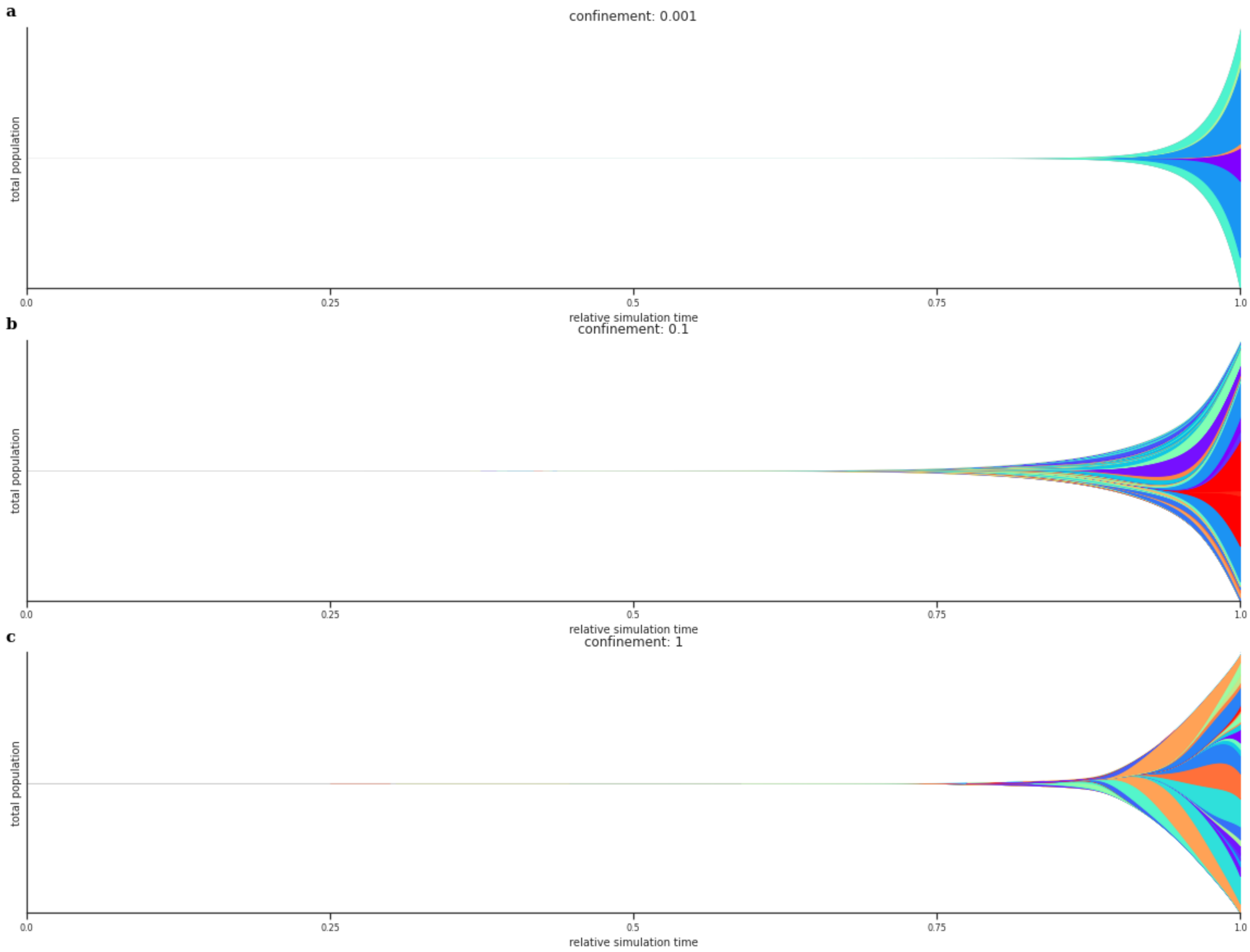


Figure 4: Three exemplary fish plots for no, intermediate and high confinement. Same as Fig. 3a-c but populations are shown in their absolute and not relative values.

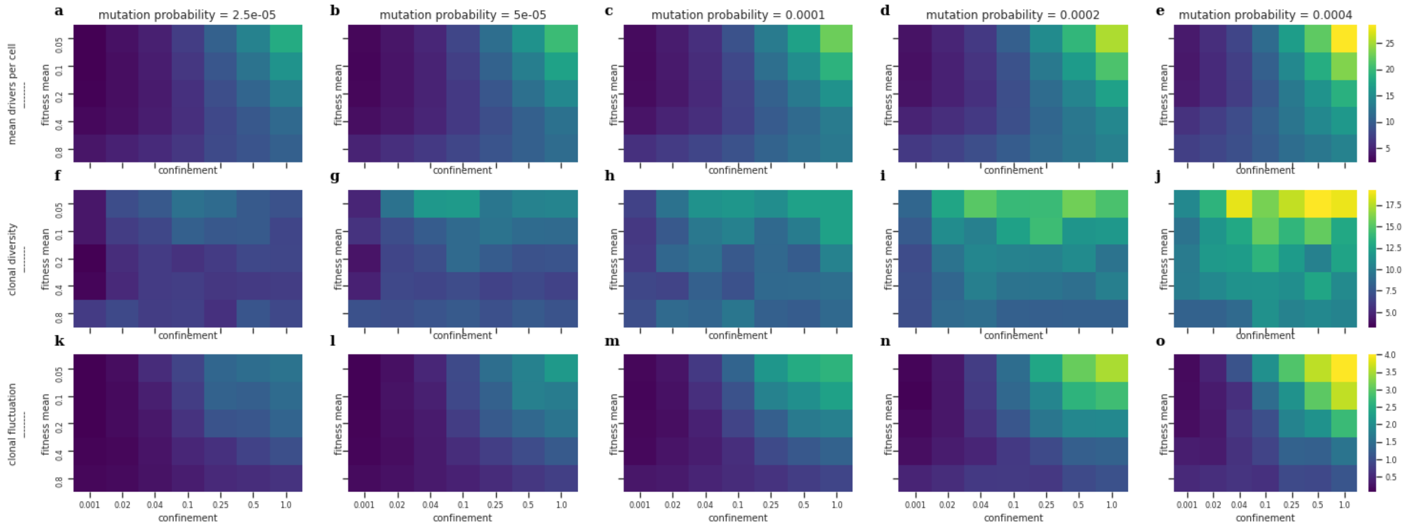


Figure 5: Full range of parameters tested in section 3.1. We see that all of the metrics are affected in non-trivial ways by the three parameters. At the same time, the general trend visible in Fig. 3 and Supp. Fig. 7 is visible as well.

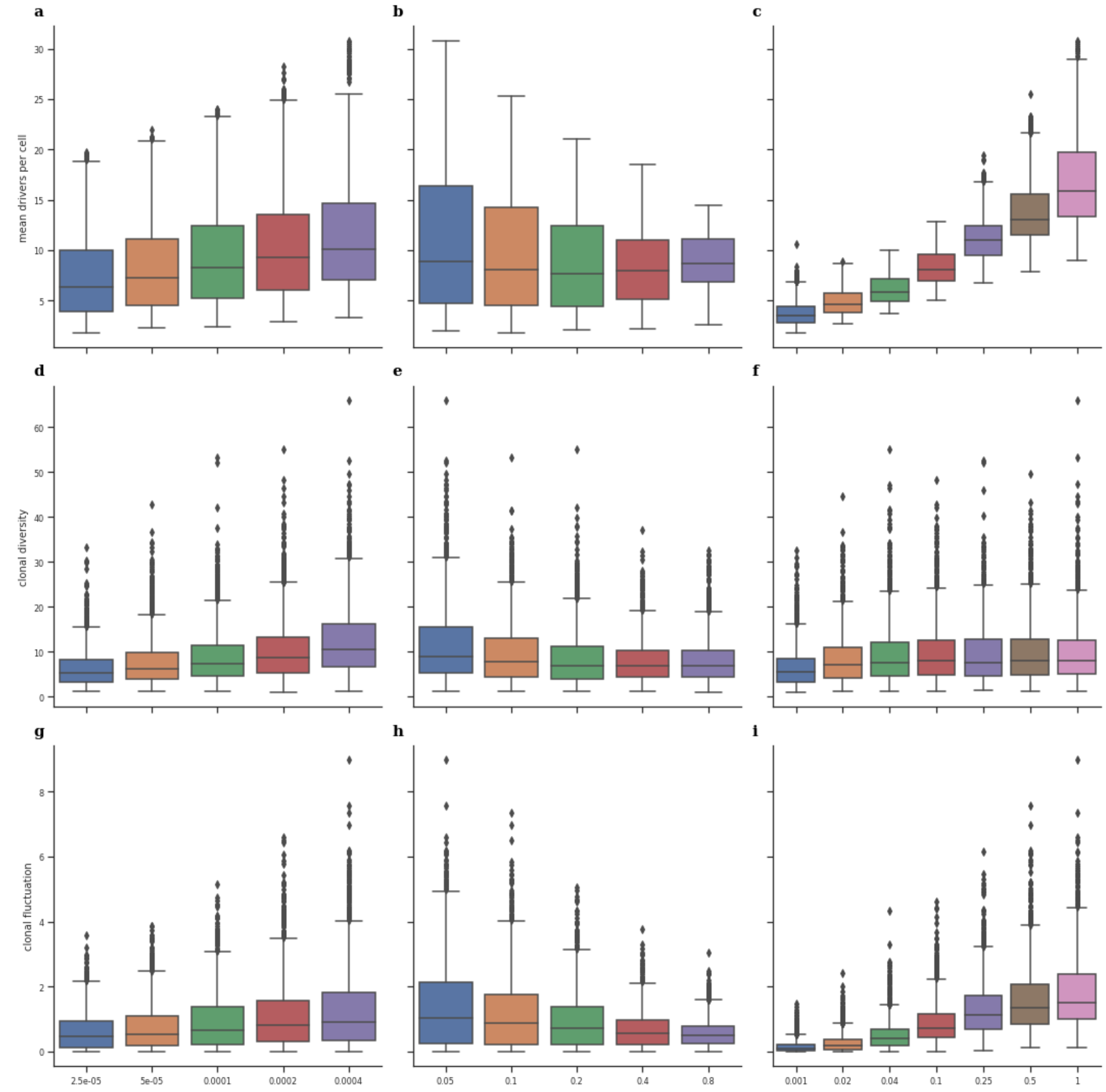


Figure 6: The effects of individual parameters on the metrics. Here we show the marginal effects of each parameter across the full range of all other parameters. Note that due to the parameter ranges chosen, the x-axis for all panels are logarithmic. A linear relationship between the y- and the x-axis therefore hint towards a logarithmic relationship between the values.

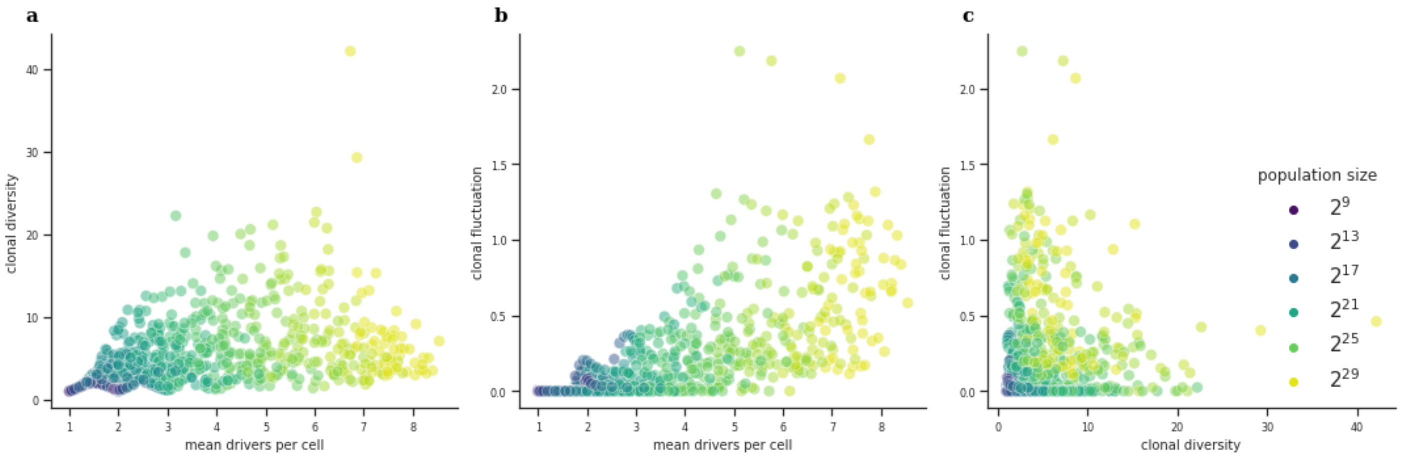


Figure 7: Pairwise evolution of the three metrics throughout the simulations for all 50 runs shown in Fig. 4a-c.

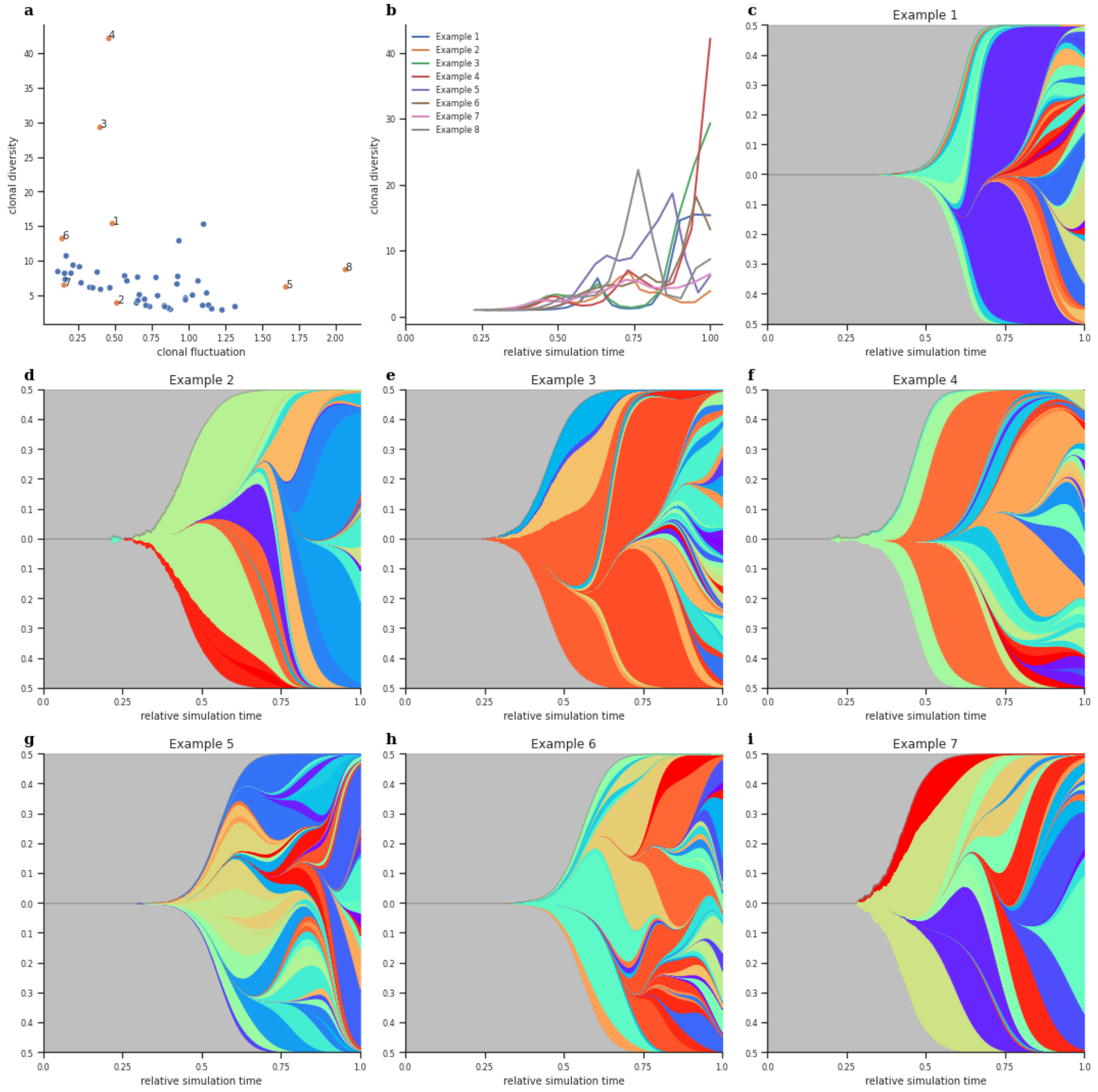


Figure 8: Examples for the high heterogeneity despite the limited amount of stochasticity and the usage of the same parameter set. **a)** For all 50 runs, the value of the final clonal diversity and the clonal fluctuation score is shown. Highlighted runs are shown in the following panels. **b)** The evolution of the clonal diversity along the simulation time for the highlighted runs. **c-i)** Fish plots for the highlighted runs. We clearly see high heterogeneity in these different simulation runs with respect to the clonal diversity and the presence/absence of bursts of diversification and clonal sweeps.

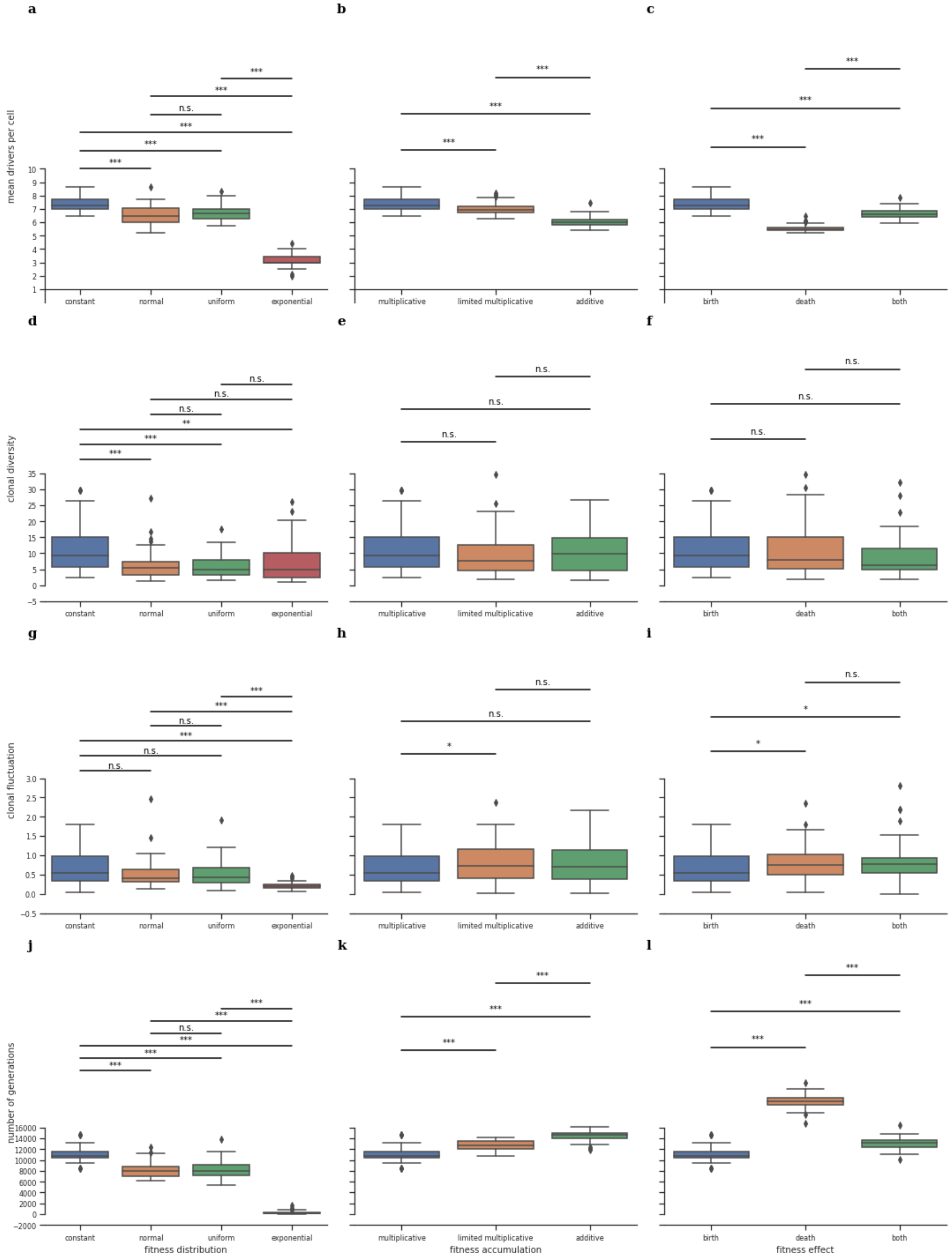


Figure 9: Full effect of the choice of the fitness distribution, accumulation method and effect as detailed in Sec. 2.4. of the main text.
